## Supplement 1 for "Identification of public submitted tick images: a neural network approach"

**SUPPLEMENTAL INFORMATION 1**

***Confusion matrix of TickIDNet predictions on the user-generated and laboratory test set.***

**S1 Table A: Confusion matrix of TickIDNet evaluated on the user-generated test set (n=1094).** The true class is the species of the tick in an image as labelled by a team of trained tick identification experts from the Midwest Center of Excellence for Vector-Borne Diseases. The predicted class is the species label assigned to the image by TickIDNet.

|  |  | **True** | | |
| --- | --- | --- | --- | --- |
|  |  | *A. americanum*  *(n = 194)* | *D. variabilis*  *(n = 625)* | *I. scapularis*  *(n = 275)* |
| **Predicted** | *A. americanum* | 151 | 24 | 7 |
|  | *D. variabilis* | 27 | 571 | 29 |
|  | *I. scapularis* | 16 | 30 | 239 |

**S1 Table B: Confusion matrix of TickIDNet evaluated on the laboratory test set (n=300).** The true class is the species of the tick in an image as labelled by TickReport; a professional tick testing and identification service. The predicted class is the species label assigned to the image by TickIDNet

|  |  | **True** | | |
| --- | --- | --- | --- | --- |
|  |  | *A. americanum*  *(n = 100)* | *D. variabilis*  *(n = 100)* | *I. scapularis*  *(n = 100)* |
| **Predicted** | *A. americanum* | 94 | 12 | 1 |
|  | *D. variabilis* | 3 | 82 | 0 |
|  | *I. scapularis* | 3 | 6 | 99 |
