## Supplement 2 for "Identification of public submitted tick images: a neural network approach"

**SUPPLEMENTAL INFORMATION 2**

**Multivariable model selection and estimates**

Evaluating the relationship between correct predictions by TickIDNet, and tick characteristics and Relative Tick Size (RTS, pixels).

**S2 Table A: Model selection.** The full model included tick characteristics (species, sex, blood feeding status) and image quality (Relative tick size [RTS]). Males and females of the same species can appear very similar (e.g., *Dermacentor variabilis*) or different (e.g., *Amblyomma americanum*) therefor we hypothesized that the relationship between sex and correct identification would not be similar for each species and we included the interaction between species and sex in our full model. Next we assessed if reduced models had a lower AIC or an AIC within 2 of the full model. As expected the full model had the lowest AIC and report on this in the manuscript, see Table S2 B for parameter estimates.

| **Model** | **Variables** | **AIC** | **deltaAIC** |
| --- | --- | --- | --- |
| **Full:** | Species * Sex + Species + Sex + Blood feeding status + RTS | 545.23 |  |
| **SBR:** | Species + Blood feeding status + RTS | 547.18 | 1.95 |
| **SSxBR:** | Species + Sex + Blood feeding status + RTS | 547.91 | 2.68 |
| **SR:** | Species + RTS | 556.04 | 10.81 |
| **BR:** | Blood feeding status + RTS | 564.66 | 19.43 |
| **R:** | RTS (univariable model) | 570.11 | 24.88 |

**S2 Table B: Model estimates for the full model.** The dataset consisted of 937 tick images; pseudo R^2^ 0.086, Area Under the Curve 0.739, Hosmer and Lemshow goodness of fit test P<.001. RTS was divided by 10; the beta reflects the change per step (i.e., 10 pixels). DV: *Dermacentor variabilis,* IS: *Ixodes scapularis*, AA: *Amblyomma americanum,* SE: Standard Error, Z: z-value, P: p-value, aOR: adjusted Odds Ratio, CI: Confidence Interval.

| **Variable** |  | **Beta** | **SE** | **Z** | **P** | **aOR** | **(95%CI)** |
| --- | --- | --- | --- | --- | --- | --- | --- |
| (Intercept) |  | 2.323 | 0.375 | 6.199 | <.001 | 10.21 | (5.06, 22.11) |
| Species | DV v. IS | -0.291 | 0.370 | -0.788 | 0.431 | 0.75 | (0.35, 1.52) |
|  | AA v. IS | -1.390 | 0.430 | -3.230 | **0.001** | 0.25 | (0.11, 0.57) |
| Sex | Male v. Female | -1.581 | 0.570 | -2.775 | **0.006** | 0.21 | (0.07, 0.66) |
| Feeding status | Fed v. Unfed | -1.161 | 0.304 | -3.813 | **<.001** | 0.31 | (0.17, 0.58) |
| RTS | (10 pixel step) | 0.126 | 0.034 | 3.720 | **<.001** | 1.13 | (1.07, 1.22) |
| Interaction Species * Sex | DV : Male | 1.749 | 0.657 | 2.663 | **0.008** | 5.75 | (1.53, 20.54) |
|  | AA : Male | 1.147 | 0.714 | 1.606 | **0.108** | 3.15 | (0.75, 12.62) |

**S2 Table C: The relationship between species and sex based on the full model, including percentage correct prediction.** The dataset consisted of 937 tick images. Adults of known sex, feeding status and species were included in the multivariable regression. The adjusted odds ratio (aOR) for correct identification accounts for image quality and feeding status (Suppl. Table 2B). Adjusted Odds Ratio illustrate how each species and sex category’s odds of correct prediction compares to *I. scapularis* females odds (reference value), CI: Confidence Interval.

| **Species** | **Female** | | | | **Male** | | | |
| --- | --- | --- | --- | --- | --- | --- | --- | --- |
|  | Correct | % | aOR | (95%CI) | Correct | % | aOR | (95%CI) |
| *I. scapularis* | 167/180 | 92.8 | 1 | Ref. level | 28/34 | 82.4 | 0.21 | (0.07, 0.66) |
| *D. variabilis* | 294/319 | 92.2 | 0.75 | (0.35, 1.51) | 238/253 | 94.1 | 0.88 | (0.04, 20.45) |
| *A. americanum* | 74/90 | 82.2 | 0.25 | (0.11, 0.58) | 46/58 | 79.3 | 0.16 | (0.01, 4.82) |

**S2 Table D: Model estimates for the SBR model.** The dataset consisted of 937 tick images; pseudo R^2^ 0.073, Area Under the Curve 0.723, Hosmer and Lemshow goodness of fit test P=0.178. RTS was divided by 10; the beta reflects the change per step (i.e., 10 pixels). DV: *Dermacentor variabilis,* IS: *Ixodes scapularis*, AA: *Amblyomma americanum,* SE: Standard Error, Z: z-value, P: p-value, aOR: adjusted Odds Ratio, CI: Confidence Interval.

| **Variable** |  | **Beta** | **SE** | **Z** | **P** | **aOR** | **(95%CI)** |
| --- | --- | --- | --- | --- | --- | --- | --- |
| (Intercept) |  | 1.992 | 0.326 | 3.116 | <.001 | 7.33 | (3.95, 14.22) |
| Species | DV v. IS | 0.111 | 0.304 | 0.365 | 0.715 | 1.12 | (0.60, 2.00) |
|  | AA v. IS | -1.194 | 0.345 | -3.462 | **<.001** | 0.31 | (0.21, 0.65) |
| Feeding status | Fed v. Unfed | -1.014 | 0.293 | -3.458 | **<.001** | 0.36 | (0.21, 0.65) |
| RTS | (10 pixel step) | 0.118 | 0.033 | 3.567 | **<.001** | 1.13 | (1.06, 1.21) |
