## Supplement 3 for "Identification of public submitted tick images: a neural network approach"

**SUPPLEMENTAL INFORMATION 3**

***Effect of image resizing on human performance***

During the image pre-processing, all images were resized from their original resolution to 224x224 pixel RGB images (see “Cropping” section). To determine whether this resizing could constrain TickIDNet’s performance we sought to quantify its impact on human performance. We asked two experts at the MCEVBD to independently score two separate, randomly sampled datasets containing 250 user-generated images each.

1) Original resolution images (no information lost)
2) Resized 224x224x3 center-cropped images (information lost)

For each dataset, we compared the inter-rater reliability between the two experts using Kohen’s Kappa (κ) measure of agreement (Table 3) that ranges from 0 (agreement equivalent to chance) to 1 (perfect agreement) (40). We hypothesized that there will be less agreement on the downscaled dataset (1) than on the original resolution dataset (2) because the downscaled images are likely more difficult to identify correctly. Notably, we did not compare accuracy because the truth labels were generated by these same experts and the physical specimens were not received for identification under a microscope.

**S3 Table: An interpretation of the Kappa statistic from Viera and Garett (40).**

| **Kappa** | **Agreement** |
| --- | --- |
| < 0 | Less than chance agreement |
| 0.01–0.20 | Slight agreement |
| 0.21–0.40 | Fair agreement |
| 0.41–0.60 | Moderate agreement |
| 0.61–0.80 | Substantial agreement |
| 0.81-0.99 | Almost perfect agreement |

The agreement between the two experts on the original resolution dataset (1) was κ = 0.9432 and the agreement on the resized image set (2) was κ = 0.8501. While S1 Table notes that both of these scores represent a near-perfect inter-rater agreement (40), the agreement on the resized images (2) was notably lower. This suggests that even human experts like the TickSpotters in Kopsco et al. (18) would likely perform slightly worse on the resized 224x224 images (seen by the network) relative to the original-resolution images. TickIDNet, just like trained experts, is constrained by image quality.
